# Enhancing HIPEC for Ovarian Cancer using Adjunctive Biomaterials

**DOI:** 10.1101/2025.10.26.684645

**Authors:** Jianling Bi, Emily Witt, Arielle B. Cafi, Fan Shu, Megan McGovern, Sri Naga Swetha Tunuguntla, Kyle R. Balk, Lillian Boge, Qi Wang, Juan Du, Ian C. Sutton, Colin J. Reis, Jacob M. Spreng, Erin J. Elizalde, Uwajachukwumma A. Uzomah, Giovanni Traverso, Leo E. Otterbein, James D. Byrne

## Abstract

Ovarian cancer is one of the most lethal gynecological malignancies, with high mortality rates primarily due to late-stage diagnoses and extensive peritoneal metastases. Despite improvements in surgical and chemotherapeutic treatments, the prognosis for advanced ovarian cancer remains poor, highlighting the urgent need for innovative therapeutic approaches. Hyperthermic intraperitoneal chemotherapy (HIPEC) has emerged as a promising treatment, delivering heated chemotherapeutic agents directly into the peritoneal cavity post-cytoreductive surgery. However, HIPEC adoption is limited by three critical complications: suboptimal therapeutic efficacy in resistant tumors; abdominal adhesion formation; and systemic toxicity, including cisplatin-induced nephrotoxicity. This study investigates carbon monoxide gas-entrapping materials (CO-GEMs) as a novel multifunctional adjunctive therapy to address the three HIPEC limitations simultaneously. CO-GEMs effectively encapsulate and deliver carbon monoxide, leveraging the differential effects of CO in cancerous where it has been shown to reduce tumor burden. In ovarian cancer models, CO-GEMs significantly enhanced cisplatin efficacy, reducing the metastatic tumor burden by 46.6% through the downregulation of drug resistance pathways, including the IL-17, TNF, and NF-κB pathways; ECM receptor interaction, and VEGF signaling. CO-GEMs also prevented peritoneal adhesion formation by suppressing inflammatory cell infiltration and collagen deposition, with a significant reduction in adhesion severity scores. Additionally, enteral CO-GEMs provided significant nephroprotection against cisplatin-induced acute kidney injury, as demonstrated by reduced blood urea nitrogen levels. CO-GEMs represent a promising innovation that simultaneously improves HIPEC therapeutic efficacy, prevents surgical complications, and reduces systemic toxicity. This multifunctional approach addresses multiple clinical limitations of HIPEC, potentially transforming treatment outcomes for patients with advanced ovarian cancer through an enhanced therapeutic index and improved safety profile.

## 1. Introduction

Ovarian cancer is one of the most lethal gynecological malignancies, ranking as the fifth leading cause of cancer-related deaths among women globally [1]. Its high mortality rate is attributed primarily to late-stage diagnoses, with most cases presenting as advanced-stage disease that is typically characterized by extensive peritoneal metastases [2]. Despite advancements in surgical and chemotherapeutic interventions, the prognosis for stage III ovarian cancer is poor, with 10-year survival rates hovering around 13%–25% [3–5]. These challenges underscore the urgent need for innovative therapeutic approaches to enhance treatment efficacy and improve patient outcomes.

Hyperthermic intraperitoneal chemotherapy (HIPEC) has emerged as a promising therapeutic modality for advanced ovarian cancer [2, 6–17]. This localized treatment involves the administration of heated chemotherapeutic agents directly into the peritoneal cavity immediately following cytoreductive surgery, thereby maximizing drug delivery to micrometastatic lesions while minimizing systemic exposure. While HIPEC has demonstrated efficacy in reducing tumor burden and extending progression-free survival in stage III ovarian cancer, its broader adoption is limited by significant complications. Three primary limitations of HIPEC are suboptimal therapeutic efficacy in chemotherapy-resistant tumors, peritoneal adhesion formation, and systemic toxicity (particularly cisplatin-induced nephrotoxicity) [18, 19]. Up to 20-30% of ovarian cancers demonstrate a lack of sensitivity to cisplatin-based chemotherapy at first line, resulting in suboptimal therapeutic efficacy [20–22]. Peritoneal adhesions represent a particularly challenging complication, arising when surgical trauma triggers inflammatory cascades involving TNF, IL-1β, and IL-6, leading to fibroblast activation and collagen deposition [23, 24]. Systemic toxicity from HIPEC is considerable, given that between 60-80% of patients experience grade 2+ toxicity following cytoreductive surgery with HIPEC, including 20-35% of patients experiencing grade 2+ cisplatin-induced nephrotoxicity [25, 26]. Together, these complications often limit the broader adoption and utility of HIPEC in clinical practice.

Recent advances in therapeutic gas delivery present a promising approach to address these HIPEC limitations [27–32]. Carbon monoxide (CO), an endogenous signaling molecule with pleiotropic biological effects, exhibits differential activity in cancerous versus normal tissues [33, 34]. CO possesses three key therapeutic properties relevant to HIPEC enhancement: (1) anti-cancer effects through metabolic disruption and chemotherapy sensitization; (2) anti-inflammatory activity that can prevent adhesion formation; and (3) cytoprotective effects that mitigate off-target organ toxicity. In particular, CO is known to suppress NF-κB-dependent cytokine production, reduce oxidative stress through HO-1/Nrf2 activation, and inhibit TGF-β/Smad pathways that drive fibroblast-to-myofibroblast transition [34–37]. Traditionally, CO has been delivered via inhalation or CO-releasing molecules, both of which are limited by titratability and carrier-associated toxicity [38]. Leveraging techniques that were originally developed in molecular gastronomy, we have created gas-entrapping materials (GEMs) composed of engineered biocompatible polysaccharide hydrocolloids [29, 30]. GEMs physically entrap therapeutic gas within a three-dimensional structure through a foaming process and facilitate controlled gas release [27–30].

This study evaluated carbon monoxide gas-entrapping materials (CO-GEMs) as a novel adjunctive therapy for cisplatin-based HIPEC in stage III ovarian cancer (Figure 1). We investigated the therapeutic effects of carbon monoxide in combination with cisplatin in ovarian cancer cell lines and assessed CO-GEM delivery systems in animal models to validate our approach. Our objectives were to (1) enhance the therapeutic efficacy of HIPEC against metastatic disease, (2) prevent peritoneal adhesion formation, and (3) reduce cisplatin-induced nephrotoxicity. The CO-GEMs leverage advanced biomaterial design principles to facilitate local and systemic gas delivery, improve tumor sensitization to cisplatin, prevent adhesions, and mitigate systemic toxicities. By integrating innovative biomaterials into existing clinical protocols, this approach has the potential to redefine management strategies for advanced ovarian cancer, ultimately improving patient outcomes.

**Figure 1.**
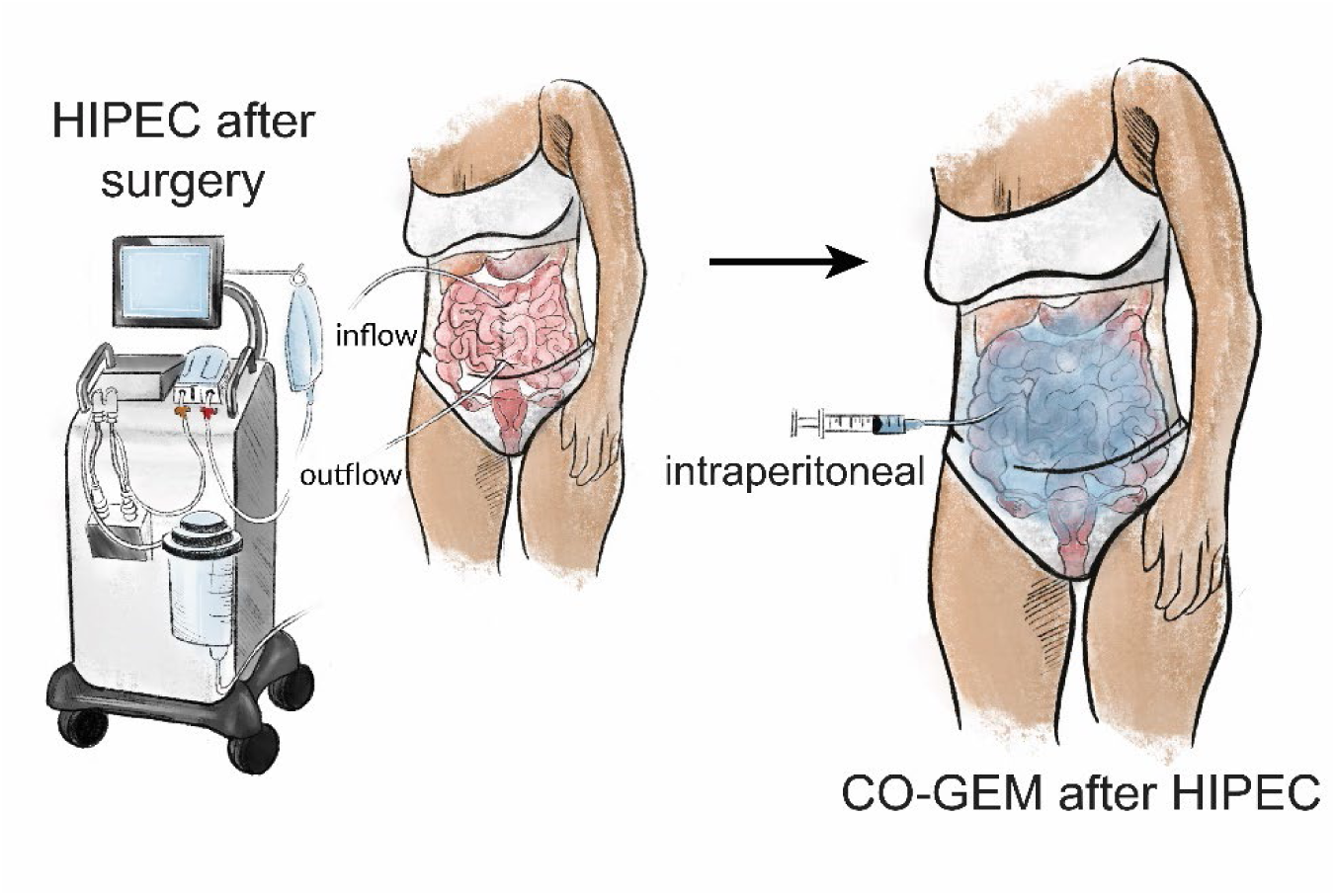
Schematic illustrating the process used to administer CO-GEM to modulate the therapeutic index of hyperthermic intraperitoneal chemotherapy (HIPEC) through intraperitoneal administration after completion of HIPEC.

## 2. Results

### 2.1. CO-GEMs are easily administered

The CO-GEMs were fabricated using a modified whipping siphon pressurized with CO to 200 pounds per square inch (psi) (**Figure 2A**). The CO-GEMs comprising of xanthan gum had the appearance of an opaque foam consisting of gas bubbles comprising 99.2% CO. The stability of the foam and the kinetics of CO release exhibited a direct, proportional relationship to the xanthan gum concentration, with higher concentrations yielding greater stability and slower release (**Figure 2B–D**). The CO-GEMs displayed viscoelastic solid-like behavior, with the storage modulus (G′) consistently surpassing the loss modulus (G″) across all tested formulations. Notably, the storage modulus increased proportionally with a higher concentration of xanthan gum (**Figure 2E**). Furthermore, each formulation demonstrated pronounced shear-thinning properties, indicating suitability for injection (**Figure 2F**). Among all the CO-GEM formulations, the 2.0 wt % xanthan gum foam exhibited a desirable balance of injectability and stability: it quickly transitioned from liquid-like flow under high shear strain to solid-like stability at lower shear strain. Consequently, this formulation was selected for subsequent evaluation in small animal models. The injectability of CO-GEMs highlights their suitability for integration into standard HIPEC protocols, facilitating efficient and reproducible application after intraperitoneal therapeutic procedures.

**Figure 2.**
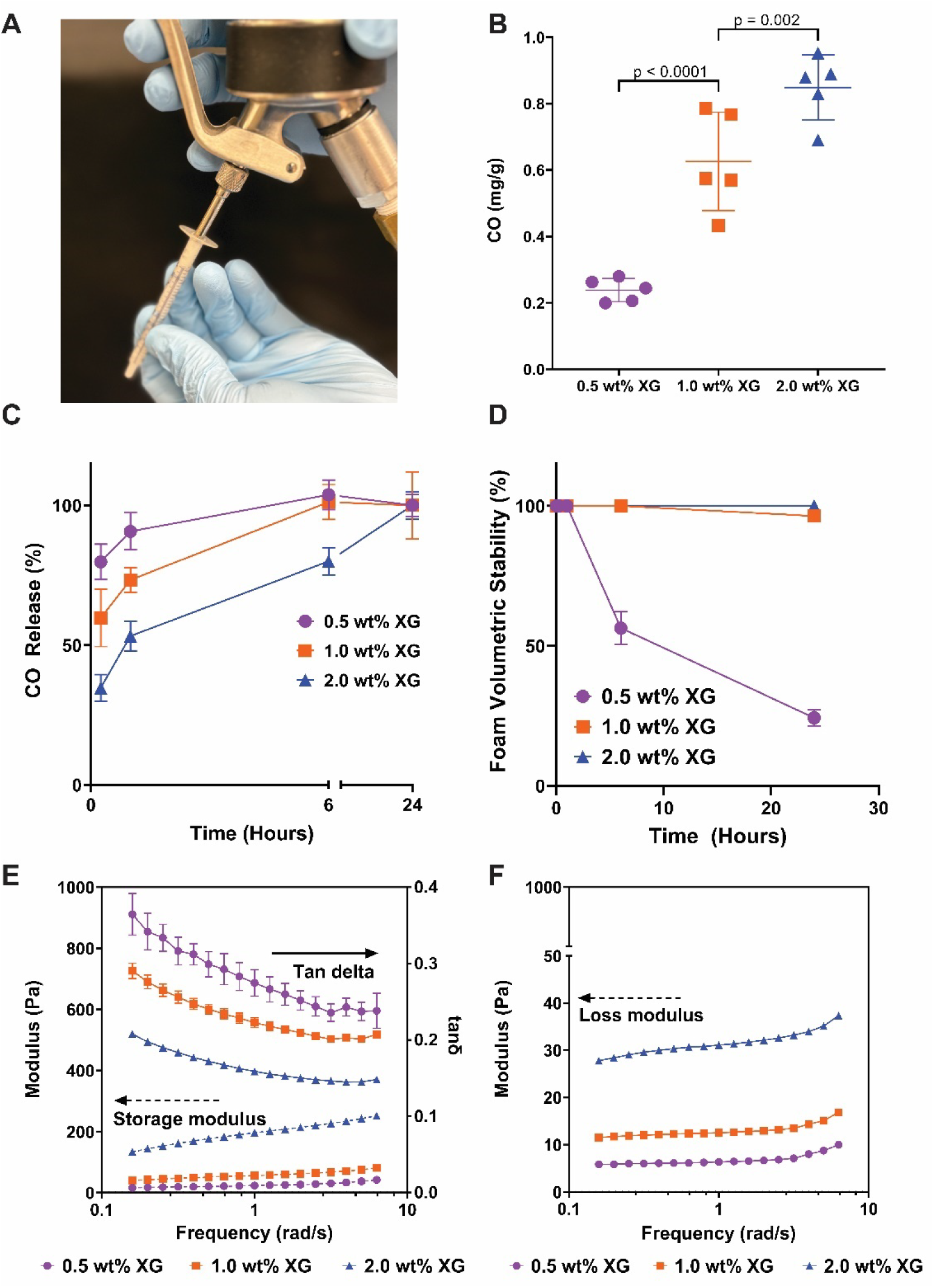
CO-GEMs contain a high concentration of CO and can be injected easily. (A) A pressurized whipping siphon shown generating the CO-GEM into a 1-mL syringe. (B) Concentration of CO in CO-GEMs at different concentrations of xanthan gum (n = 5/group). P-value was determined by one-way ANOVA. (C) CO release kinetics from different CO-GEMs at different concentrations of xanthan gum (n = 3/group). (D) Foam volumetric stability of CO-GEMs at different concentrations of xanthan gum (n = 3/group). (E) Storage modulus increased proportionally with higher concentration of xanthan gum (n = 3/group). (F) Loss modulus of CO-GEMs increased with xanthan gum concentration, with all formulations maintaining elastic-dominant behavior (G’ > G’’) across the frequency range (n = 3/group).

### 2.2. Intraperitoneal administration of CO-GEMs achieves elevated COHb %

Achieving therapeutic carbon monoxide levels is critical for CO-GEM efficacy, as subtherapeutic concentrations fail to provide anti-tumor benefits while excessive levels can cause toxicity, making carboxyhemoglobin percentage (COHb %) monitoring essential for safe and effective dosing. The pharmacokinetics of CO from the CO-GEMs were evaluated in C57BL/6 mice after intraperitoneal delivery (2.5 g/kg). The COHb %, a measure of the percentage of hemoglobin that is bound to CO, reached a maximum directly after CO-GEM administration and decreased over 6 hours (**Figure S1**). Intraperitoneal administration achieved a similar COHb % compared to both oral and rectal administration [27, 30].

### 2.3. Intraperitoneal CO-GEMs enhance cisplatin efficacy and reduce metastatic burden in ovarian cancer

Given the significant rates of cisplatin resistance in ovarian cancer, we hypothesized that carbon monoxide co-treatment would overcome therapeutic limitations by sensitizing cancer cells to cisplatin and amplifying cytotoxic effects. We first investigated whether CO could enhance the cytotoxic effects of cisplatin in 5 human ovarian cancer cell lines (OVCAR3, OVCAR4, OV-7, CAOV3, SKOV3). Cytotoxicity was assessed using an alamarBlue assay. We found that treatment of these cell lines concurrently with cisplatin plus CO (250 ppm) (cisplatin + CO) significantly reduced the half-maximal inhibitory concentration (IC₅₀) compared to cisplatin alone (**Table S1**). This enhanced cytotoxic effect was accompanied by increased markers of apoptosis (cleaved poly (ADP-ribose) polymerase (PARP)) and DNA damage (phosphorylated γ-H2AX) (**Figure S2**). To understand the mechanism underlying increased cisplatin sensitivity for ovarian cancer cells, we performed RNA sequencing on OVCAR3 cells exposed to 250 ppm CO. The analysis revealed significant downregulation of multiple pathways associated with drug resistance, including IL-17, TNF, NF-κB, ECM receptor interaction, VEGF signaling, and platinum drug resistance (**Figure S3** and **S4**). Additionally, in vitro migration of OVCAR3 cells was significantly impaired upon exposure to cisplatin + CO compared to either treatment alone (**Figure S5**).

We next evaluated the impact of CO-GEM on metastatic burden in an OVCAR3 peritoneal metastases mouse model simulating advanced-stage ovarian cancer (**Figure 3A–B**). Intraperitoneal administration of CO-GEMs following cisplatin-based HIPEC resulted in a significant decrease in metastatic burden (**Figure 3C–D**). Quantitative analysis revealed a 46.6% reduction in overall tumor volume in mice treated with HIPEC + CO-GEM compared to HIPEC + control GEM. These results demonstrate that CO-GEMs can enhance the regional therapeutic impact of cisplatin HIPEC by suppressing drug resistance pathways and modulating cytokine signaling.

**Figure 3.**
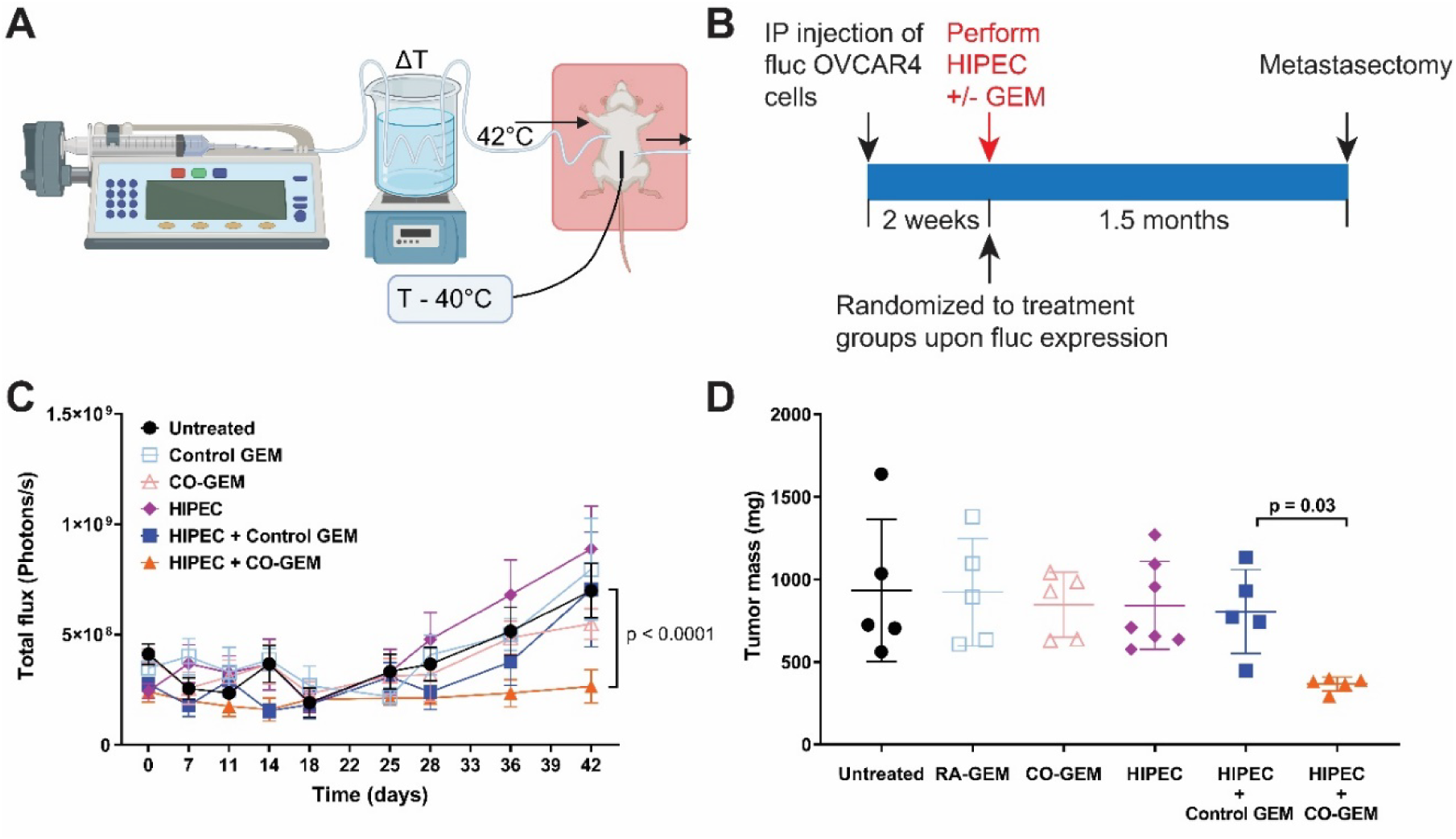
CO-GEM treatment as an adjunct to cisplatin HIPEC leads to lower metastatic burden in an ovarian cancer peritoneal metastases animal model. (A) Illustration of experimental setup for HIPEC. (B) Timeline for the experimental design where the firefly luciferase (fluc) expressing OVCAR4 cells were injected into athymic (nude) female mice, and 2 weeks later, were evaluated for fluc expression via bioluminescence. Mice with luciferase expression were randomized into equal groups and then exposed to HIPEC + CO-GEM or Control (Room Air (RA)) GEM. Metastasectomy was performed approximately 1.5 months after the HIPEC treatments. (C) Total flux for bioluminescence imaging of the ovarian peritoneal cancer model for the different treatment groups (n = 5-7/group). P-value was determined by linear regression. (D) Tumor masses post-metastasectomy of the ovarian peritoneal cancer mouse model. P-value was determined by one-way ANOVA.

### 2.4. CO-GEMs prevent peritoneal adhesion formation through anti-inflammatory mechanisms

Given that peritoneal adhesions represent a major complication of HIPEC, resulting from procedure-induced inflammatory cascades [26], and that CO exhibits established anti-inflammatory properties [35], we hypothesized that CO-GEMs could prevent adhesion formation. Using an established mouse adhesion model involving cecal abrasion and abdominal wall injury [39], we found that intraperitoneal CO-GEMs significantly reduced adhesion incidence compared to controls, with treated mice showing a 3-fold reduction in adhesion severity scores (scoring system in Methods) (**Figure 4A**). Histological evaluation using α-smooth muscle actin (α-SMA), CD68, and Masson trichrome staining revealed that CO-GEM treatment significantly reduced inflammatory cell infiltration (CD68), activated myofibroblasts (α-SMA), and collagen deposition (Masson trichrome) compared to control GEMs (**Figure 4B**). This adhesion prevention is mechanistically consistent with the ability of CO to suppress NF-κB-dependent cytokine production, reduce oxidative stress through HO-1/Nrf2 activation, and inhibit TGF-β/Smad pathways that drive fibroblast-to-myofibroblast transition [40, 41].

**Figure 4.**
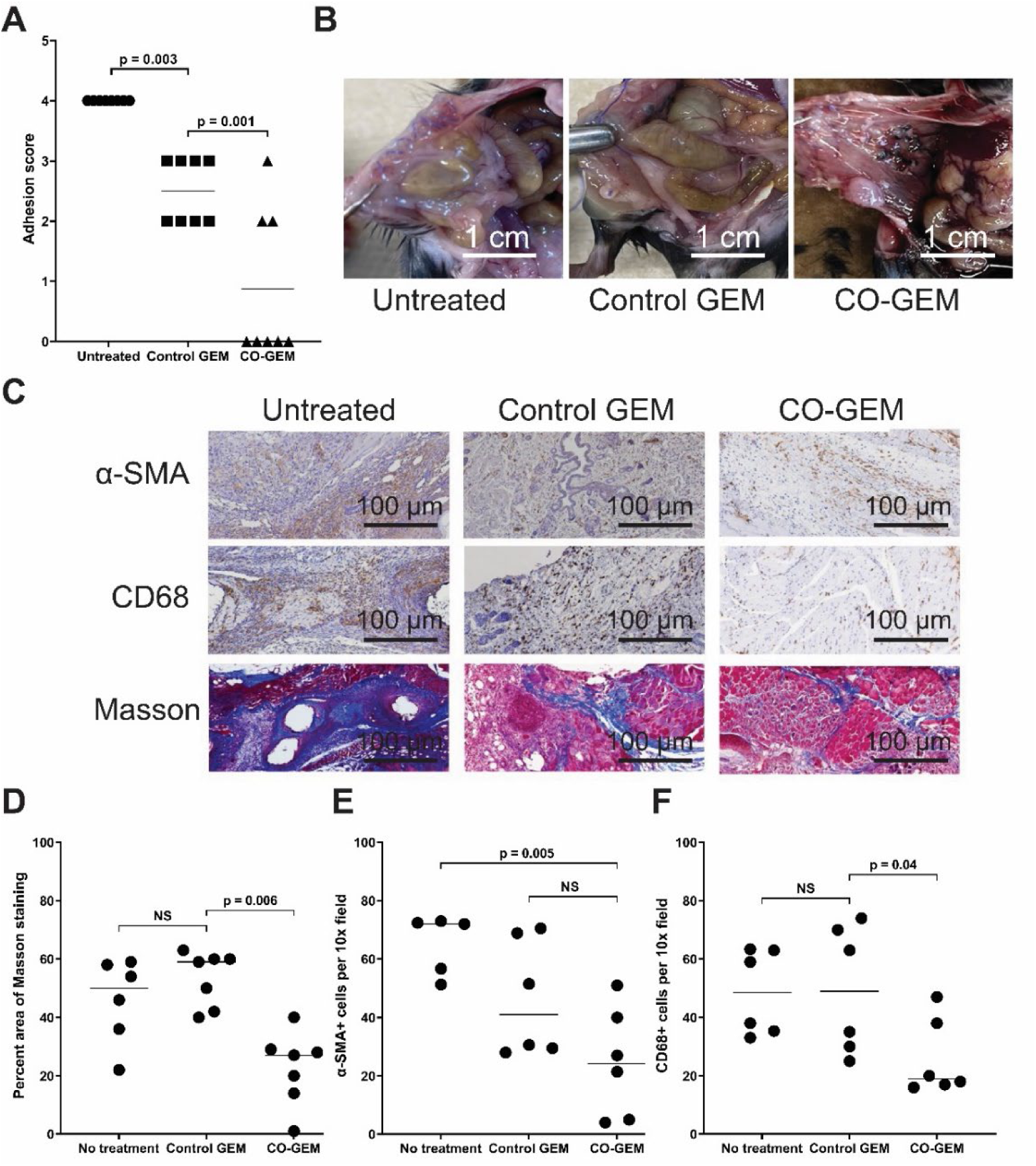
CO-GEMs treatment reduced abdominal adhesions in a mouse adhesion model (C57BL/6 female mice). P-value was determined by one-way ANOVA. (A) Adhesion scores for mice treated with no treatment, control GEM (2.5 mg/kg), or CO-GEMs (2.5 mg/kg) (n = 8/group). P-value determined by one-way ANOVA. (B) Representative images of the anterior abdominal wall of mice treated with no treatment, control GEM, or CO-GEMs. (C) Representative histologic imaging of the anterior abdominal wall. (D) Percent area of positive staining in 10x field for Masson trichome and (E) α-SMA+ cells per 10x field, and (F) CD68+ cells per 10x field (n = 5/group). P-value was determined by one-way ANOVA.

To further elucidate the anti-inflammatory mechanisms by which CO-GEMs mitigate adhesion formation, we examined the effects of CO on macrophage oxidative stress and inflammatory signaling. Using murine bone marrow–derived macrophages (BMDMs), CO treatment significantly reduced intracellular reactive oxygen species (ROS) levels as assessed by DCFH-DA fluorescence microscopy and flow cytometry (**Figure S6**). Under lipopolysaccharide (LPS) stimulation, CO continued to suppress ROS production, indicating a broad antioxidant effect (**Figure S7**). At the transcriptional level, CO markedly downregulated pro-inflammatory gene expression, including inducible nitric oxide synthase (NOS2) and arginase 1 (ARG1), consistent with attenuation of macrophage activation (**Figure S8**). Collectively, these data demonstrate that CO-GEMs modulate macrophage oxidative and inflammatory responses, likely contributing to reduced fibroinflammatory remodeling and peritoneal adhesion formation observed *in vivo*.

### 2.5. CO-GEMs prevent cisplatin-induced acute kidney injury in mice

Systemic toxicity, particularly cisplatin-induced acute kidney injury (AKI), represents a major dose-limiting factor for HIPEC. Based on the established cytoprotective properties of CO in normal tissues [33], we investigated whether CO-GEMs could mitigate cisplatin nephrotoxicity while preserving anti-cancer efficacy. First, we confirmed that low-dose CO was cytoprotective in HK-2 renal epithelial cells during cisplatin exposure (**Figure S9**). We then determined that enteral administration of CO-GEMs in a cisplatin-induced AKI mouse model provided significant nephroprotection, reducing blood urea nitrogen (BUN) levels by 58% compared to the control group (**Figure 5A**).

**Figure 5.**
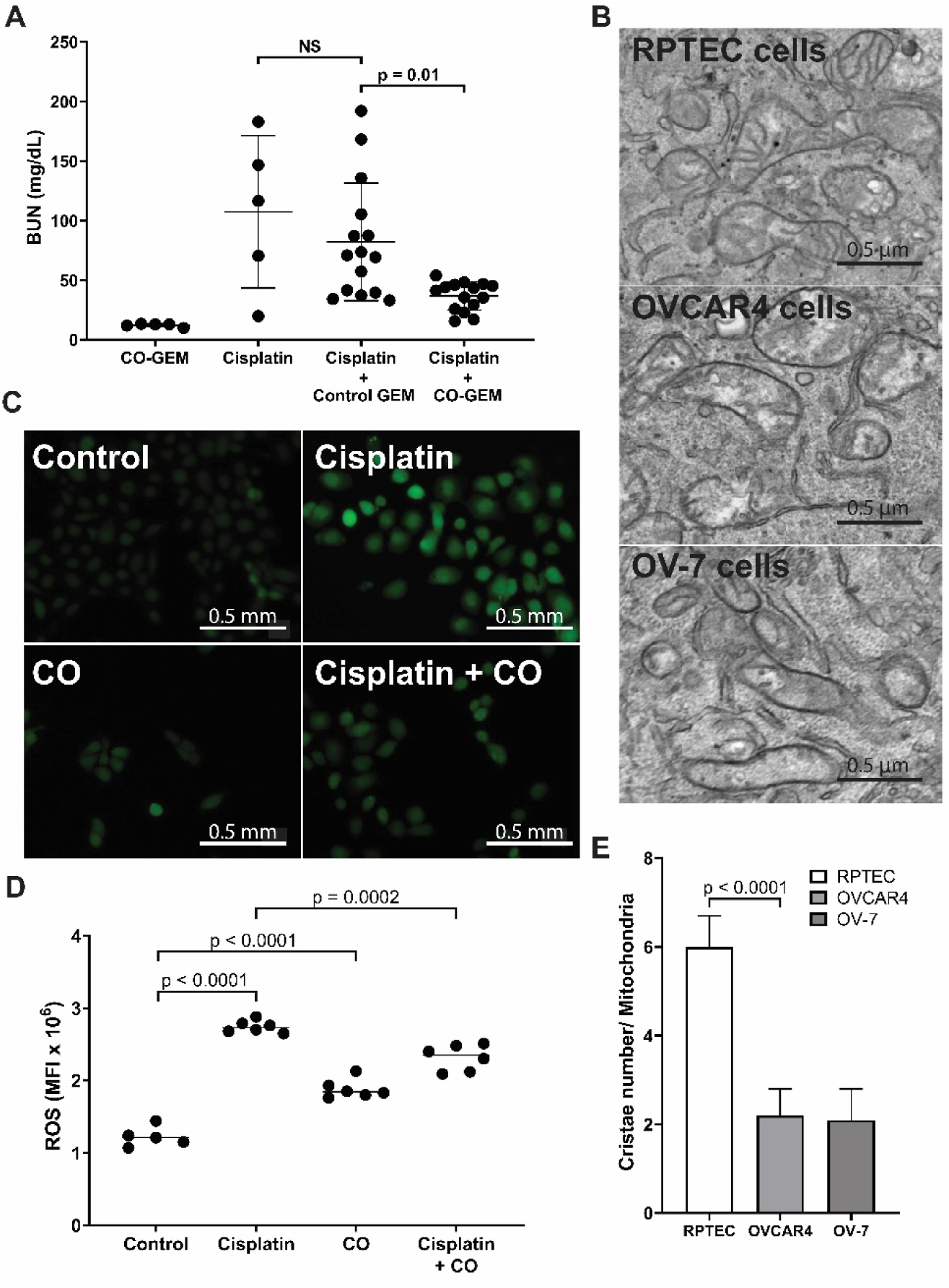
CO-GEMs protect against cisplatin-induced kidney injury. (A) Serum blood urea nitrogen (BUN) levels of mice treated with CO-GEM (2.5 mg/kg, n = 5), cisplatin (15 mg/kg intraperitoneal, n = 5), cisplatin + control GEM (n = 15), or cisplatin + CO-GEM (n = 15). CO-GEM significantly reduced cisplatin-induced BUN elevation (p = 0.01), whereas control GEM provided no protection (NS). P-value determined by one-way ANOVA. (B) Transmission electron micrographs of normal kidney cells (RPTEC) compared to ovarian cancer cells (OVCAR4 and OV-7), showing differential mitochondrial cristae architecture. (C) Representative fluorescence microscopy images of normal human proximal tubular cells exposed to cisplatin, CO, cisplatin + CO, or control (untreated), cells were pretreated with 250 ppm CO for 3h, then exposed to 10nM cisplatin for 24h. Cells were stained with 2’,7’-dichlorodihydrofluorescein diacetate (DCFH-DA) for ROS detection. (D) Quantification of DCFH-DA fluorescence via flow cytometry. Cisplatin significantly increased ROS levels (p < 0.0001), which was reduced by CO co-treatment (p = 0.0002). (E) Quantification of cristae number per mitochondrion in RPTEC cells versus ovarian cancer cells (OVCAR4 and OV-7) from subfigure B (n = 15/group). Normal kidney cells exhibit significantly higher cristae density (p < 0.0001). P-value determined by one-way ANOVA.

To understand the differential effects of CO on cancer versus normal cells, we compared mitochondrial profiles between ovarian cancer cells and primary renal proximal tubule epithelial cells, building on previous evidence of renal protection. Transmission electron microscopy revealed that normal primary renal proximal tubule epithelial cells (RPTEC) possessed significantly more cristae per mitochondria compared to two ovarian cancer cell lines (OVCAR4, OV-7) (**Figure 5B** and **E**). An increased density of cristae expands the inner mitochondrial membrane surface area, supporting more efficient oxidative phosphorylation and enhanced respiratory capacity in normal cells compared to cancer cells. We hypothesize that this structural difference allows normal kidney cells to better tolerate transient CO-mediated cytochrome c oxidase inhibition through redundant respiratory chain complexes, whereas cancer cells with fewer cristae become more susceptible to CO-induced metabolic disruption. Furthermore, we evaluated 2’,7’-dichlorodihydrofluorescein diacetate (DCFH-DA) staining for reactive oxygen species (ROS) in renal proximal tubule epithelial cells and found that CO co-treatment with cisplatin reduced ROS formation in these cells (**Figure 5C** and **D**). In addition, CO modulates the inflammatory gene expression within these cells, leading less IL6 and higher CCL2 mRNA transcription in renal proximal tubule epithelial cells treated with cisplatin (**Figure S10**), indicating a selective, reparative inflammatory profile.

## 3. Discussion

We investigated the effectiveness of CO-GEMs to simultaneously address three critical limitations of HIPEC treatment in ovarian cancer. CO-GEMs effectively entrap substantial volumes of CO, enabling intraperitoneal and enteral delivery of CO. We found that intraperitoneal administration of CO-GEMs following cisplatin-based HIPEC significantly reduced metastatic burden in an ovarian cancer metastasis mouse model, primarily driven by the suppression of key drug resistance-associated pathways and the modulation of cytokine signaling pathways. Additionally, CO-GEMs addressed critical limitations associated with HIPEC, notably reduced therapeutic efficacy in some patients due to reduced cisplatin sensitivity, abdominal adhesions and cisplatin-induced acute kidney injury. Collectively, these results highlight the effectiveness and versatility of CO-GEMs in enhancing therapeutic outcomes and mitigating the significant complications associated with HIPEC in the treatment of ovarian cancer.

Currently, no adjunctive strategies are clinically utilized to enhance the therapeutic index of HIPEC specifically. Existing biomaterial applications in HIPEC patients have primarily been limited to abdominal wall closure aimed at preventing hernia formation [42]. Preclinical studies have explored alternative biomaterial strategies, such as cytokine delivery systems, which demonstrated significant potential [43, 44]. Our biomaterial innovation uniquely leverages gas-entrapping materials (GEMs) to physically encapsulate CO within a polysaccharide hydrocolloid matrix, facilitating controlled and targeted gas release [27–32, 45]. This study demonstrates that CO-GEMs simultaneously address three critical limitations of HIPEC, potentially transforming clinical outcomes for patients with advanced ovarian cancer.

CO functions as a multifaceted therapeutic agent that simultaneously enhances cancer treatment efficacy while mitigating treatment-related complications through its differential effects on malignant versus normal cellular physiology. In cancer cells, CO induces metabolic exhaustion (anti-Warburg effect) and reduces key resistance factors, including glutathione and metallothionein levels [33, 46]. Our RNA sequencing data support this mechanism, showing CO-mediated downregulation of drug resistance pathways, including those involved in IL-17, TNF, NF-κB, and platinum drug resistance. CO prevents peritoneal adhesion formation by interrupting the inflammatory cascade at multiple points. Surgical trauma typically triggers acute inflammation (TNF, IL-1β, IL-6) and oxidative stress, leading to a fibrin-rich provisional matrix that matures into fibrous adhesions under persistent cytokine signaling [23, 24]. CO delivery prevents this progression through (1) immune modulation via NF-κB/p38-MAPK suppression and IL-10 promotion, (2) anti-oxidative signaling through HO-1/Nrf2 activation, (3) anti-fibrotic effects via TGF-β/Smad pathway restraint, and (4) mesothelial protection that preserves non-adhesive surfaces [40, 41]. Systemic protective effects against cisplatin toxicity appear to be related to differential mitochondrial architecture between normal versus cancer cells. The enhanced cristae density of normal kidney cells that we observed in this study provides greater respiratory capacity, enabling better tolerance of transient CO-mediated cytochrome c oxidase inhibition compared to cancer cells with compromised mitochondrial structure [47]. Indeed, CO has been shown to induce mitochondrial biogenesis [48]. Together, the enhanced therapeutic outcomes observed in our study may be attributable to the local and systemic release of CO and the retention of the therapeutic gas within the intraperitoneal cavity.

This study has several limitations, which provide critical directions for future research. First, our findings are derived from a single animal model, which limits their generalizability across diverse physiological contexts. Future research should validate these findings in additional models, including immunocompetent and orthotopic systems, to more accurately reflect clinical complexity. Second, titration studies to define an optimal therapeutic window for CO administration were not performed. Systematic dose-response investigations are crucial for determining the minimal effective and maximal tolerated doses, thereby guiding safe clinical translation. Third, the current CO-GEM formulation relies exclusively on xanthan gum as the hydrocolloid matrix. Future exploration of alternative polysaccharide-based materials may enhance tunable release kinetics, biocompatibility, and formulation stability. Nevertheless, the robust therapeutic outcomes observed provide a strong foundation for the continued development and diversification of the CO-GEM platform. Finally, while we noted downregulation of several key inflammatory pathways, including IL-17, TNF, and NF-κB, comprehensive transcriptomic and mechanistic studies remain necessary to fully elucidate the broad biological effects of CO and identify additional therapeutic targets.

Overall, CO-GEMs represent a promising adjunctive approach to enhance the therapeutic index of HIPEC for ovarian cancer and potentially other intraperitoneal malignancies. The distinctive properties of GEMs facilitate a scalable, modular strategy for localized delivery of therapeutic gases, potentially extending beyond CO to include other biologically active gases. The simplicity of the formulation, utilization of polysaccharide hydrocolloids, and ambient pressure manufacturing process support its future clinical translation and integration into existing intraperitoneal administration techniques [49]. Importantly, CO-GEMs directly address a critical clinical need by improving the efficacy of chemotherapy while simultaneously reducing cisplatin-associated toxicities without altering the chemotherapeutic agent itself. Additionally, addressing regulatory considerations, such as dose standardization, formulation reproducibility, and manufacturing process scalability, will be essential for advancing CO-GEMs toward clinical application. Ultimately, these efforts may enable the incorporation of CO-GEMs into standard HIPEC protocols (**Table S2**) [2, 6–17], offering advanced ovarian cancer patients a safer and more effective therapeutic option.

## 4. Materials and methods Formulation development

CO-GEMs were prepared by first dissolving xanthan gum (Sigma-Aldrich) in ultrapure water under continuous stirring, while heating to 100 °C. Once fully dissolved, the solution was allowed to cool to room temperature and was subsequently degassed using a vacuum chamber. The degassed solution was then transferred into a modified whipping siphon, which was preloaded and pressurized to 200 psi with either pure CO or medical-grade air (for controls), depending on the experimental condition.

### Material characterization

CO-GEMs were visually inspected, followed by microscopic evaluation using an EVOS microscope (10X). Foam stability was assessed by transferring 100 mL of foam into a 100-mL graduated cylinder. Samples were incubated at 37 °C and observed at 0 minutes, 15 minutes, 1 hour, 6 hours, and 24 hours. At each time point, the volumes of foam and liquid were recorded to assess stability over time. CO content was analyzed by gas chromatography with a thermal conductivity detector (Agilent). CO gas from three foam formulations containing xanthan gum at 0.5%, 1.0%, and 2.0% by weight was quantified using a Hewlett Packard 6890 Series gas chromatograph (GC). Before gas quantification, GC vials were subjected to multiple purges to eliminate background gases. Foams were then injected into screw-cap vials and allowed to rest for either 15 minutes, 1 hour, 6 hours, or 24 hours to assess time-dependent CO release. For each time point, five replicates were prepared. From each vial, 500 µL of headspace gas was extracted and injected into the GC. CO concentrations were calculated based on calibration curves created with 99.5% carbon dioxide. Rheological measurements were carried out using a Kinexus Ultra+ rheometer (Malvern Panalytical, Malvern, UK). For each formulation, frequency sweeps ranging from 0.1 to 20.0 Hz (0.65 to 130.00 rad/s) were conducted at a strain of 1% under controlled temperature conditions (37 °C). Experiments were performed in triplicate using roughened stainless steel parallel plates with a diameter of 20 mm. Foam gels were gently placed between the plates and were compressed to a gap height of 1.5 mm, applying less than 0.05 N of normal force.

### In vitro studies—cell viability assays

To evaluate the impact of exogenous CO on cisplatin cytotoxicity, we performed cytotoxicity assays on multiple cell lines. For the control condition, OVCAR3, OVCAR4, OV-7, CAOV3, and SKOV3 cells were plated in 96-well plates at a density of 10,000 cells per well and allowed to adhere for 24 hours. Cells were then exposed to a range of cisplatin concentrations (0.1 µg/mL – 10 µg/mL) for 72 hours in a standard incubator with 5% CO₂ in air. For the CO-treated condition, the same cell lines were plated under identical conditions, but cisplatin treatment was performed within a sealed exposure chamber (StemCell Technologies) containing 250 ppm CO blended with 5% CO₂ in air for 72 hours. In a separate set of experiments, HK-2 and RPTEC cells were seeded at the same density and incubated for 24 hours, followed by a 3-hour exposure to 250 ppm CO in 5% CO₂-balanced air. After this pre-exposure, the cells were treated with varying concentrations of cisplatin for an additional 72 hours. Cell viability was determined using the alamarBlue cell proliferation reagent (ThermoFisher Scientific) according to the manufacturer’s instructions. Absorbance was measured with a microplate reader (Bio-Rad Laboratories, Hercules, CA) at an excitation wavelength of 560 nm and an emission wavelength of 590 nm. All assays were performed with three technical replicates, and viability values were normalized to those of untreated controls, which were defined as 100%.

### In vitro studies—western blots

Cells were harvested and lysed in RIPA buffer (sc-24948; Santa Cruz Biotechnology, Dallas, TX). Protein samples (40–60 µg) were loaded onto either 10% or 12% SDS– polyacrylamide gels, separated by electrophoresis, and transferred to nitrocellulose membranes (PALL Corporation, Port Washington, NY). Membranes were blocked with 5% non-fat dry milk and incubated overnight at 4 °C with the appropriate primary antibody. The following primary antibodies were used (Cell Signaling Technology, Danvers, MA): anti-cleaved PARP (1:1000, #95696); anti–phospho-γ-H2AX (1:1000, #9718); anti–phospho-ERK (1:1000, #4370); anti-phospho-c-jun-Ser73 (1:1000, #3270); anti-c-jun(1:1000, #9165); anti-phospho-Ampkα-Thr172 (1:1000, #2535), anti-Ampkα(1:1000, #2532); anti-α-tubulin(1:1000, #2144); and anti-ERK (1:1000, #4695). HO-1(1:500, sc-10789), β-actin (1:10000, sc-47778; Santa Cruz Biotechnology, Dallas, TX) was used as a loading control. Protein bands were visualized using the Bio-Rad ChemiDoc imaging system, and densitometric analysis was performed with Bio-Rad Image Lab software (Bio-Rad Laboratories, Redmond, WA).

### In vitro studies—RNA sequencing

OVCAR3 cells, OVCAR4 cells, and BMDMs were exposed to 250 ppm CO for 48 hours prior to RNA isolation. Total RNA was extracted using TRIzol Reagent (Simgen, Hangzhou, China) following the manufacturer’s instructions, and residual genomic DNA was eliminated with DNase I (Takara Bio, Shiga, Japan). RNA quality was evaluated using an Agilent 2100 Bioanalyzer (Agilent, Palo Alto, CA), and RNA quantity was quantified with a NanoDrop ND-2000 (Thermo Scientific, Waltham, MA). Only samples meeting the following criteria were used for library construction: Optical Density (OD) 260/280 between 1.8 and 2.2; OD 260/230 ≥ 2.0; RNA integrity number (RIN) ≥ 6.5; 28S:18S ratio ≥ 1.0; and yield >1 µg.

RNA-seq libraries were prepared from 1 µg of total RNA using the TruSeq™ RNA Sample Preparation Kit (Illumina, San Diego, CA). Poly(A)+ mRNA was enriched with oligo(dT) beads and fragmented, followed by synthesis of double-stranded cDNA using the SuperScript IV kit (Invitrogen, Waltham, MA). Subsequent steps (end repair, phosphorylation, and A-tailing) were performed according to Illumina’s protocol. Libraries (∼300 bp) were size-selected on 2% Low Range Ultra Agarose, amplified by PCR with Phusion DNA polymerase (NEB; 15 cycles), quantified with a TBS380 fluorometer, and sequenced on an Illumina NovaSeq 6000 Sequencing System (paired-end, 2 × 150 bp).

Raw paired-end reads were adapter-trimmed and quality-filtered using SeqPrep and Sickle (default parameters). Filtered reads were aligned to the reference genome in orientation mode with HISAT2 [50] and assembled with StringTie [51] using a reference-guided strategy. Transcript and gene abundance were estimated with RSE [52] and expressed as transcripts per million (TPM). Differential expression was determined with DESeq2 [53] /DEGseq [54] /edgeR [55], applying significance thresholds of |log2 fold change| > 1 and Q ≤ 0.05 (DESeq2/edgeR) or Q ≤ 0.001 (DEGseq). Gene Ontology (GO) and KEGG pathway enrichment analyses were performed using GOATOOLS and KOBAS [52], respectively, with a Bonferroni-adjusted p-value of ≤0.05. Alternative splicing events were identified with rMATS [56] (including isoforms corresponding to reference annotations or containing novel splice junctions) and categorized as exon skipping or inclusion, alternative 5′ or 3′ splice sites, or intron retention.

### In vitro studies—migration assays

OVCAR3, OVCAR4, and OV-7 cells were serum-starved overnight. A suspension containing 3 × 10⁴ cells in 150 µl of serum-free medium was placed into the upper chamber of a 6.5-mm Transwell insert with an 8-µm pore polycarbonate membrane (3422; Corning, Glendale, AZ). The lower chamber contained 600 µl of medium supplemented with 10% FBS as a chemoattractant. Cells were incubated for 16 hours in the presence or absence of 250 ppm CO. Following incubation, non-migrated cells on the upper membrane surface were removed, and cells on the underside of the membrane were fixed with 4% paraformaldehyde, stained with 0.2% crystal violet for 30 minutes, and counted in five randomly selected fields at 20× magnification.

### In vitro studies—BMDM polarization

Bone marrow-derived macrophages (BMDMs) were isolated from the femurs and tibias of 6–8-week-old C57BL/6 mice. Bone marrow cells were flushed, subjected to red blood cell lysis and filtration, and cultured in complete DMEM supplemented with M-CSF (20 ng/mL) for 5 days to obtain unpolarized M0 macrophages, with medium replacement on day 3. On day 6, the culture medium was pretreated with 250 ppm CO and then applied to BMDMs for 3 h, followed by stimulation with LPS (50 ng/mL) and IFN-γ (10 ng/mL) for 24 h. Gene expression was analyzed by qPCR. The primers used are listed: 18S rRNA: 5′-TCAACTTCGATGGTAGTCGCCGT-3′, and 5′-TCCTTGGATGTGGTAGCCGTTCT-3′; Nos2: 5′- GAGACAGGGAAGTCTGAAGCAC-3′ and 5′-CCAGCAGTAGTTGCTCCTCTTC-3′; Arg1: 5′-CATTGGCTTGCGAGACGTAGAC-3′ and 5′-GCTGAAGGTCTCTTCCATCACC-3′.

### In vitro studies—fluorescence microscopy and flow cytometry

BMDMs were obtained from mouse femurs and cultured with 20 ng/mL M-CSF for 5 days. Cells were pre-treated with 250 ppm CO for 3 hours, followed by 1μg/mL or 0.1μg/mL LPS stimulation for 1 h. Cells were then stained with DCFH-DA for 1 hour. ROS levels were measured using green fluorescence and FITC on a Nikon inverted microscope ECLIPSE Ts2R-FL and a BD FACSCelesta flow cytometer.

### In vitro studies—electron microscopy

RPTEC, OVCAR4, and OV-7 cells were cultured on 18-mm glass coverslips placed in 12-well plates for 48 hours. Cells were rinsed with 2 mL of DPBS and fixed overnight at 4 °C in a solution containing 2% formaldehyde and 2.5% glutaraldehyde in 0.1 M cacodylate buffer (pH 7.4). Post-fixation was performed using 1% osmium tetroxide for 1 hour. Samples were dehydrated through a graded ethanol series (50%, 75%, 95%, and 100%) and embedded in Epon 12 resin (Ted Pella, Redding, CA). Ultrathin sections (∼70 nm) were prepared using an ultramicrotome and were subsequently stained with uranyl acetate followed by lead citrate. Imaging was performed using a Hitachi HT7800 transmission electron microscope (Tokyo, Japan) at the University of Iowa Central Microscopy Research Facility.

### Animal studies

All animal experiments were performed under a protocol approved by the University of Iowa Institutional Animal Care and Use Committee (IACUC #1092429-008). Female C57BL/6J mice (6–8 weeks old) and CD-1 mice (4 weeks old) were obtained from the Jackson Laboratory (Bar Harbor, ME) and housed in the University of Iowa animal care facility. Animals were given a 72-hour acclimation period before study initiation, were maintained on a 12-hour light/dark cycle and were provided food and water ad libitum.

### Animal studies—HIPEC study

C57BL/6 mice bearing established ovarian peritoneal carcinomatosis were used to evaluate the feasibility and morbidity of chemotherapy administered by HIPEC. Luciferase-expressing OVCAR-3 cells (6 × 10⁶) were injected intraperitoneally 21 days prior to treatment. Tumor engraftment and progression were verified using bioluminescence imaging, and the mice were randomized to different treatment groups. The mice were monitored using bioluminescence imaging on day 19 post-inoculation (2 days before HIPEC) and weekly thereafter until sacrifice. Each mouse underwent a single HIPEC procedure and was monitored daily for 14 days post-treatment to assess survival and potential treatment-related morbidity. HIPEC was performed as follows: A heating mattress set to 35 °C was determined to be effective for maintaining normothermia during the procedure. Anesthesia was induced using 4% isoflurane in oxygen (2 L/min) delivered via an inhalation chamber. Analgesia was administered preoperatively by subcutaneous injection of buprenorphine (0.5 mL). The abdominal cavity was accessed through multiple small incisions to accommodate the placement of inflow and outflow catheters, enabling controlled intraperitoneal perfusion. Throughout the HIPEC procedure, perfusate temperature and flow rate were closely monitored and adjusted as needed to maintain consistent thermal delivery. After completion of HIPEC, the cisplatin solution was removed from the peritoneal cavity using the perfusion catheter. Subsequently, 200 µL GEMs (CO or control Room Air) were administered into the peritoneal cavity directly after completion of HIPEC. Body temperature was continuously recorded using a rectal probe to ensure physiological thermal stability during treatment.

### Animal studies—adhesion study

We followed an established protocol to generate an adhesion mouse model [39]. Mice were pre-treated with antibiotic chow (Irradiated Uniprim Diet; Inotiv, Lafayette, IN) for 1 week prior to surgery. Under isoflurane anesthesia and sterile conditions, the abdomen was shaved, disinfected, and opened via a midline laparotomy to expose the peritoneal cavity. The cecum was gently externalized and abraded with fine-grit sandpaper to induce mild serosal injury, followed by a more aggressive abrasion and placement of figure-of-eight sutures (4-0 silk) along the right abdominal wall to promote adhesion formation. Warmed saline was used for intra-abdominal irrigation, and medical starch was applied to both the cecum and sidewall to enhance adhesion development. After ensuring hemostasis, the intestines were returned to the cavity, and the abdomen was closed in two layers. Postoperatively, mice received subcutaneous fluids and were dried and bandaged before being returned to recovery. The histologic adhesion scoring system employed a 6-point scale (0-5) based on adhesion density and tissue contact area. Score 0 indicated no adhesions, while score 1 represented minimal string-like adhesions with up to 10% tissue contact. Scores 2 and 3 described progressively thicker, noncontinuous adhesions with tissue contact up to 25% and 50%, respectively. Score 4 indicated continuous adhesions with multiple contact points covering up to 75% of tissue, and score 5 represented the most severe category with dense, continuous adhesions covering 100% of adherent tissue surface area [57, 58].

### Animal studies—acute kidney injury study

CD-1 mice were housed under standard conditions with controlled temperature (18–24 °C), relative humidity (40%–70%), and a 12-hour light/dark cycle. Food and water were available ad libitum. Prior to experimentation, all mice were acclimated in a conventional housing environment for 1 week. For treatment, mice received oral gavage of 100 μL of either CO-GEMs or control room air GEMs 90 minutes prior to being administered cisplatin (15 mg/kg, intraperitoneally). At 96 hours post-cisplatin treatment, mice were euthanized for sample collection. Blood was drawn for analysis of blood urea nitrogen (BUN) levels. The left kidney was preserved in 4% phosphate-buffered formalin for histological evaluation.

### Statistical analysis

Data are displayed as individual points or as means accompanied by standard deviations. Statistical analyses and graph generation were done using GraphPad Prism 10 software (Boston, MA). We compared the treatment groups using either analysis of variance (ANOVA) or unpaired t-tests, as appropriate.

## CRediT authorship contribution statement

**Jianling Bi:** Conceptualization, Methodology, Formal Analysis, Investigation, Resources, Writing – Original Draft, Visualization, and Supervision. **Emily Witt:** Methodology, Validation, Formal Analysis, Investigation, Writing – Review & Editing. **Arielle B. Cafi:** Investigation, Writing – Review & Editing. **Fan Shu:** Investigation, Writing – Review & Editing. **Megan McGovern:** Investigation, Writing – Review & Editing. **Sri Naga Swetha Tunuguntla:** Investigation, Writing – Review & Editing. **Kyle R. Balk:** Investigation, Writing – Review & Editing. **Lillian Boge:** Investigation, Writing – Review & Editing. **Qi Wang:** Investigation. **Juan Du:** Investigation. **Ian C. Sutton:** Investigation, Writing – Review & Editing. **Colin J. Reis:** Investigation, Writing – Review & Editing. **Jacob M. Spreng:** Investigation, Writing – Review & Editing. **Erin J. Elizalde:** Investigation, Writing – Review & Editing. **Uwajachukwumma A. Uzomah:** Investigation, Writing – Review & Editing. **Giovanni Traverso:** Conceptualization, Writing – Review & Editing. **Leo E. Otterbein:** Conceptualization, Writing – Review & Editing. **James D. Byrne:** Conceptualization, Writing – Original Draft, Supervision, Project Administration, Funding Acquisition.

## Acknowledgement

We thank the team at SayoStudio for their illustration in Figure 1. The authors also thank the staff of the University of Iowa Central Microscopy Research Facility for their rapid and detailed work on the histology reported in this study. We thank Dr. Kristan Worthington for access to their rheometer. This work was funded in part by grants from the University of Iowa Hospitals and Clinics Department of Radiation Oncology; the Holden Comprehensive Cancer Center at The University of Iowa and its National Cancer Institute (NCI) Awards P30CA086862 (JDB), NCI K08CA276908 (JDB), and NIH DP2CA301081 (JDB); the American Cancer Society IRG-21-141-46 (JDB); the Prostate Cancer Foundation Young Investigator Award (JDB); the Department of Defense Prostate Cancer Program Early Investigator Award W81XWH-20-1-0225 (JDB); the Department of Defense W81XWH-16-0464 (LEO); the National Football League Players Association (LEO); the National Science Foundation 1927616 (MST); the Karl van Tassel (1925) Career Development Professorship (GT); and the Massachusetts Institute of Technology Department of Mechanical Engineering (GT). We would like to thank the Baylor College of Medicine Mouse Metabolism & Phenotyping core (NIH fund RO1DK114356 & UM1HG006348) for their analysis of our mouse plasma samples.

