## Supplemental Figures and Table for "Enhancing HIPEC for Ovarian Cancer using Adjunctive Biomaterials"

**Supplemental Information**

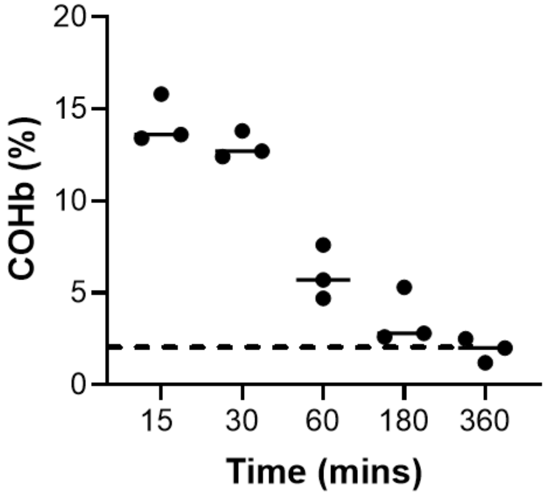


**Figure S1.** Percentage of carboxyhemoglobin (COHb %) measured for intraperitoneal administration of CO-GEMs (2.5 mg/kg) at the indicated times (C57BL/6J female mice, n = 5 per group). The dotted line represents the highest baseline COHb %.

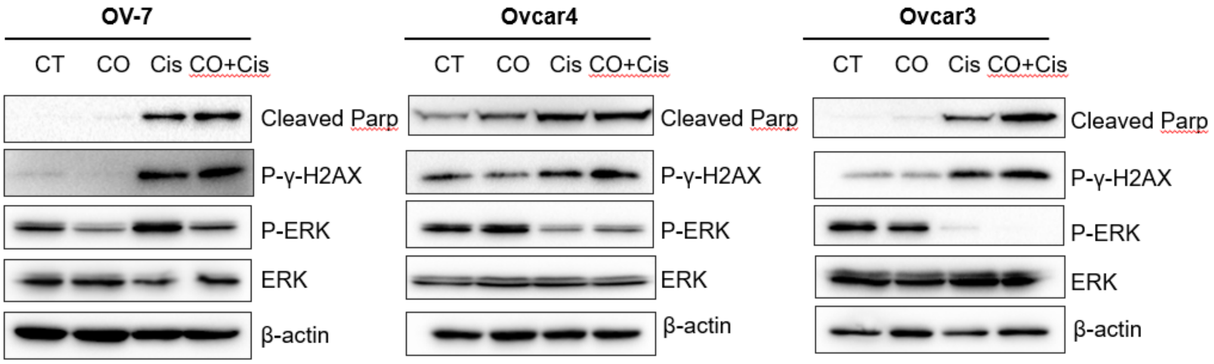


**Figure S2.** The combination of CO (250 ppm) + cisplatin increased markers of DNA damage (P-γ-H2AX) and apoptosis (Cleaved Parp) in OV-7, OVCAR4, and OVCAR3, as demonstrated via western blots. Cisplatin: 0.5 µg/ml OVCAR3; 1 µg/ml OVCAR4; and 1µg/ml OV-7 at 72 hours.

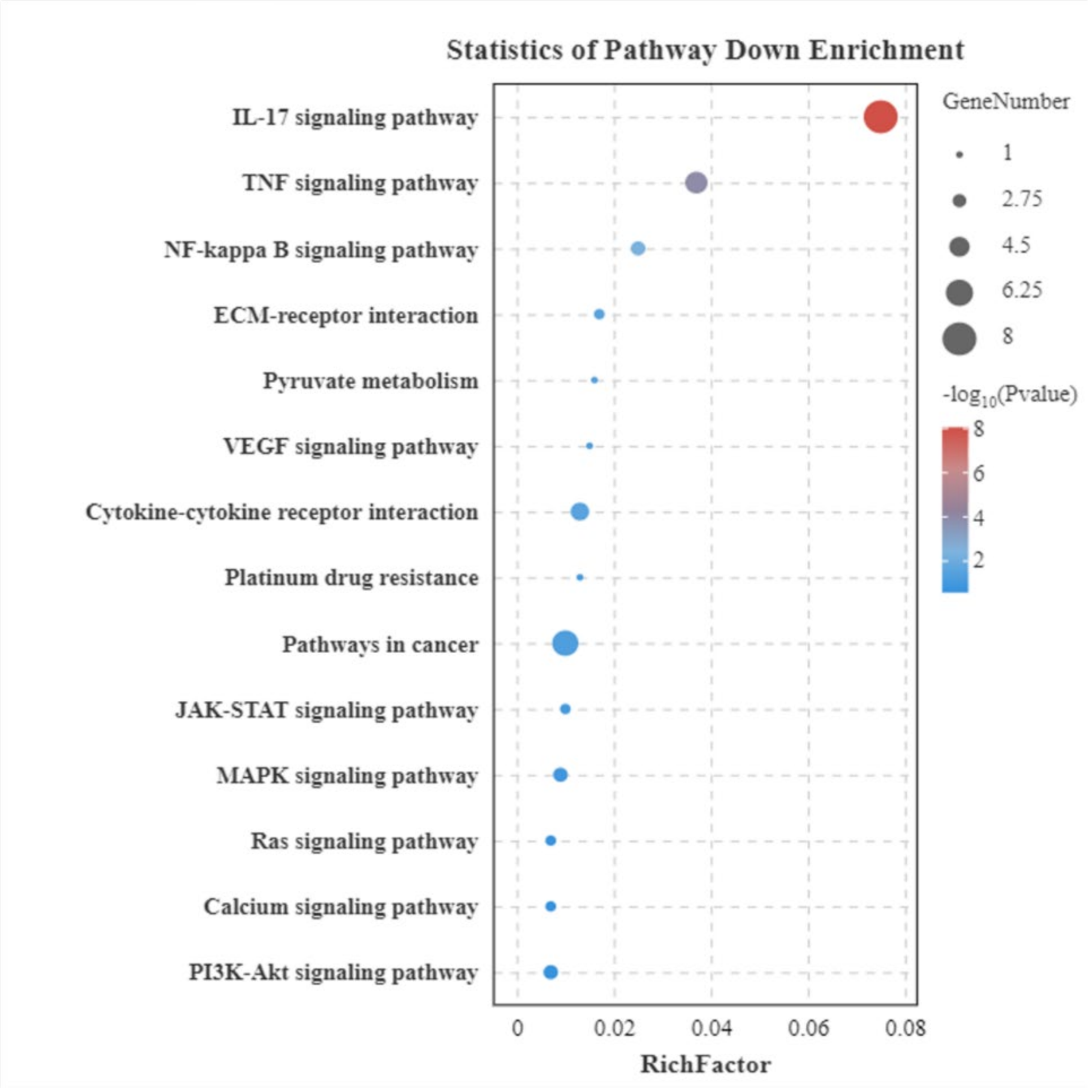


**Figure S3.** Heat maps from RNA sequencing analysis of OVCAR3 cells lines exposed to air (control) or 250 ppm of CO for 48 hours.

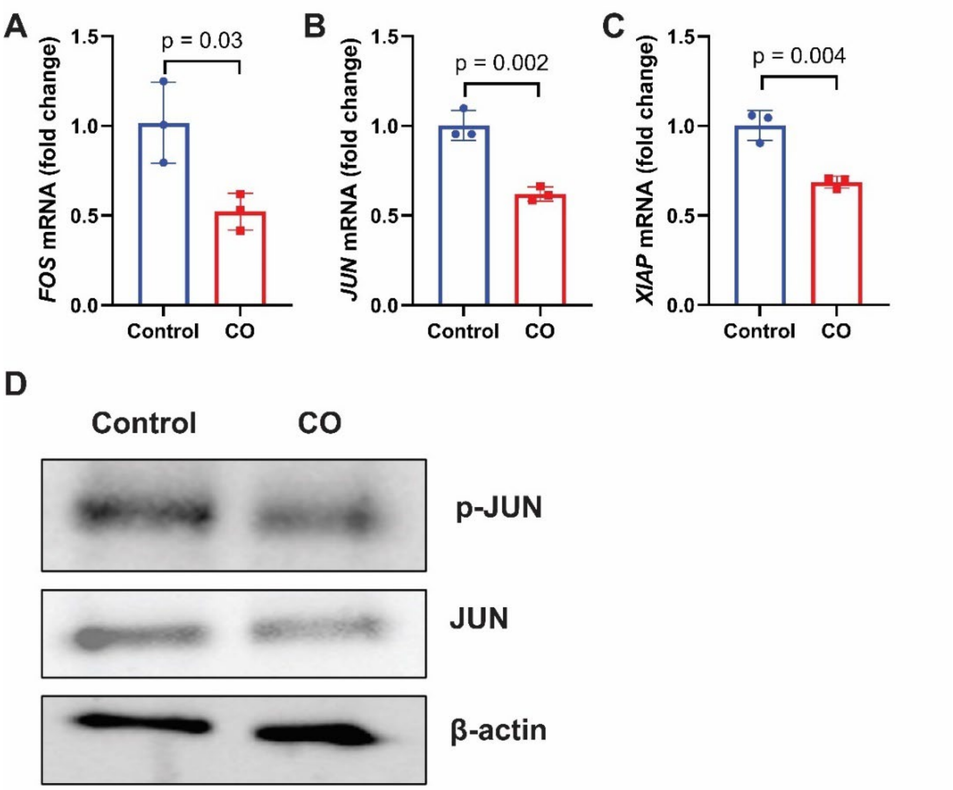


**Figure S4.** CO reduces expression of chemoresistance-associated genes. (A-C) mRNA expression of FOS (A), JUN (B), and XIAP (C) in OVCAR3 cells treated with control or 250ppm CO for 24 h. P value was determined by unpaired t-test. CO treatment significantly reduced expression of FOS ($p = 0.03$), JUN ($p = 0.002$), and XIAP ($p = 0.004$) compared to control ($n = 3$-$4$). Data represent mean ± SD fold change relative to control. (D) Representative western blot showing protein levels of phosphorylated JUN (p-JUN), total JUN, and β-actin in cells treated with control or 250ppm CO. CO treatment reduced both p-JUN and total JUN protein levels.

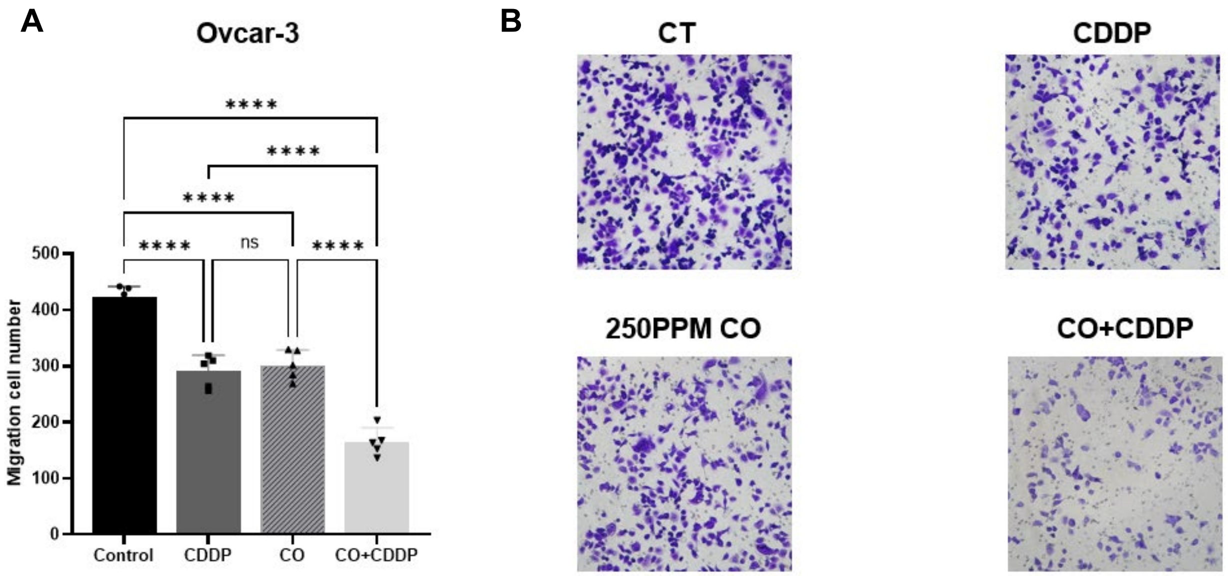


**Figure S5.** CO (250 ppm) + cisplatin (CDDP) reduces OVCAR3 cell migration. (A) Quantification of OVCAR3 cells exposed to CO + cisplatin, CO, cisplatin, or untreated control (n = 5 per group) migrating through a Transwell. P-value was determined by one-way ANOVA. (B) Crystal violet staining of the transmembrane.

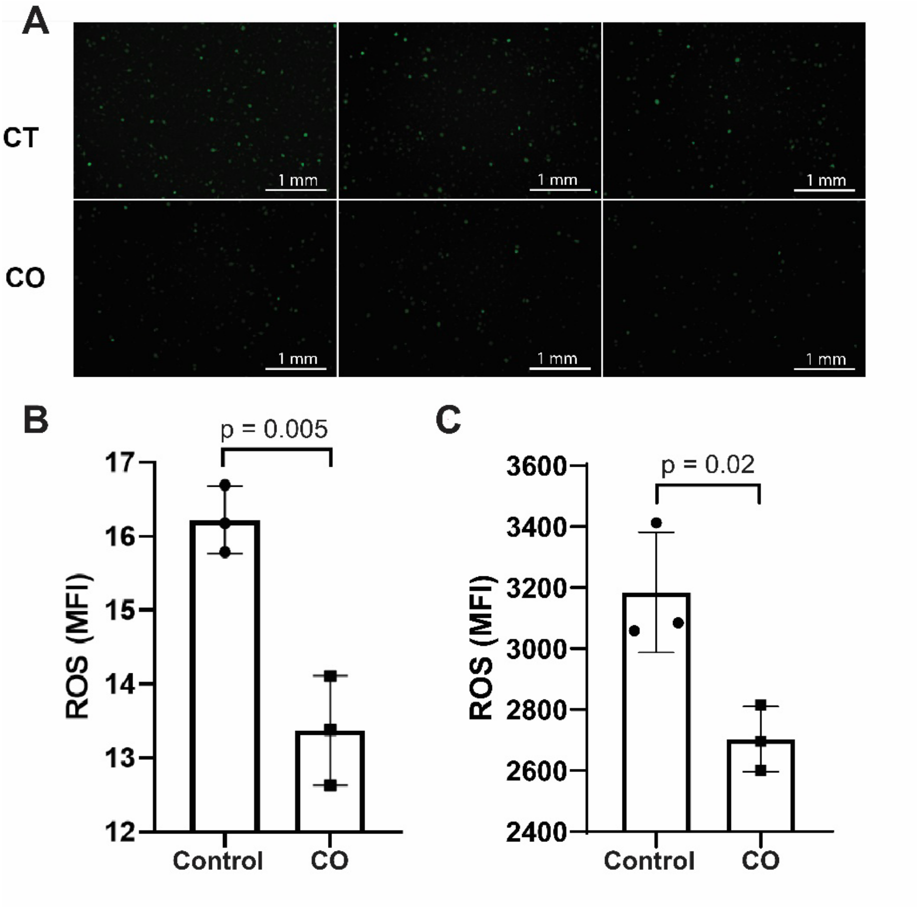


**Figure S6.** CO reduces LPS-induced reactive oxygen species in murine bone marrow-derived macrophages (BMDMs). BMDMs were pretreated with 250ppm CO for 3 hours, then exposed to 1 μg/ml LPS for 1 hour. (A) Representative fluorescence microscopy images of BMDMs treated with control (CT) or CO stained with 2',7'-dichlorodihydrofluorescein diacetate (DCFH-DA) to detect reactive oxygen species (ROS). Three technical replicates are shown per condition. Green fluorescence indicates ROS generation. (B) Quantification of ROS levels by MFI for panel A. CO treatment significantly reduced ROS compared to control ($p = 0.05$, $n = 3$). Data represent mean ± SD. P value was determined by unpaired t-test. (C) Quantification of ROS levels by mean fluorescence intensity (MFI) via flow cytometry. CO treatment significantly reduced ROS compared to control ($p = 0.002$, $n = 3$). Data represent mean ± SD. P value was determined by unpaired t-test.

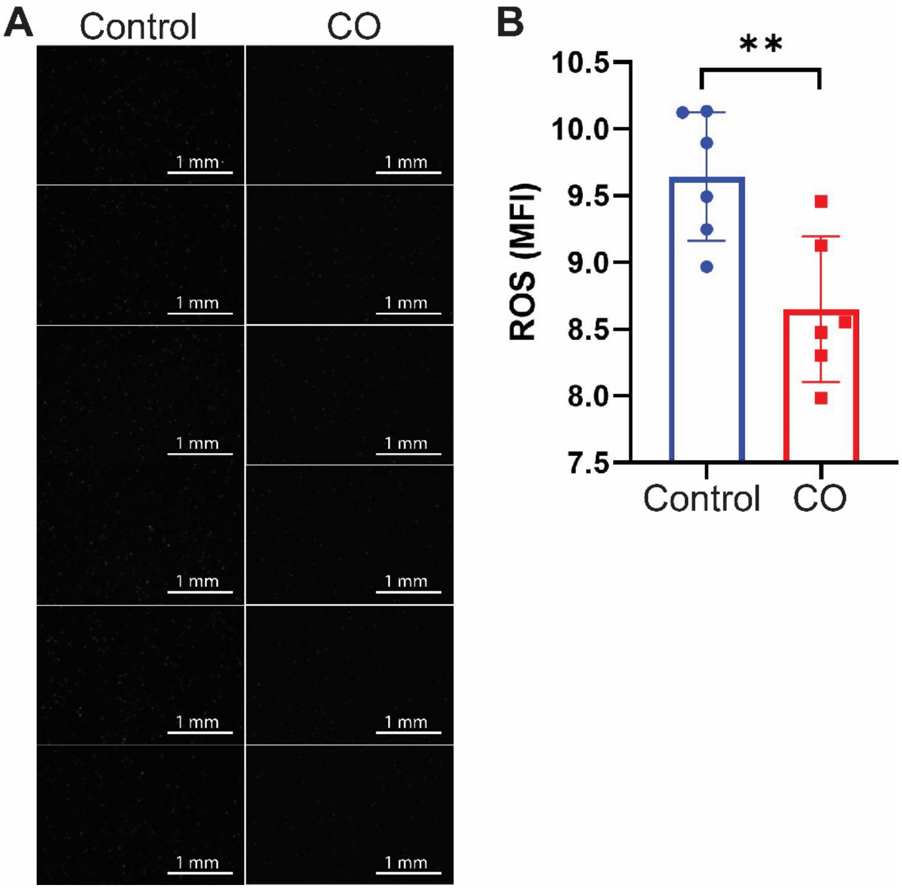


**Figure S7.** CO reduces ROS generation in BMDMs under reduced LPS stimulation. BMDMs were pretreated with 250ppm CO for 3 hours, then exposed to 0.1 μg/ml LPS for 1 hour. (A) Representative fluorescence microscopy images of BMDMs stained with 2',7'-dichlorodihydrofluorescein diacetate (DCFH-DA) to detect reactive oxygen species (ROS). Six technical replicates are shown per condition. (B) Quantification of ROS levels by mean fluorescence intensity (MFI) via flow cytometry. CO treatment significantly reduced ROS compared to control (** $p < 0.01$, $n = 6$). Data represent mean ± SD. P value was determined by unpaired t-test.

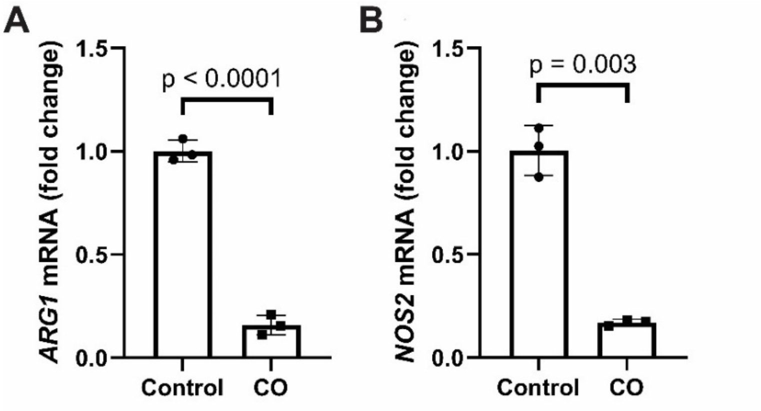


**Figure S8.** CO reduces pro-inflammatory gene expression in BMDMs. (A) mRNA expression of ARG1 (arginase 1) in BMDMs treated with control or 250 ppm CO for 24h. CO treatment significantly reduced ARG1 expression compared to control ($p < 0.0001$, n = 3). Data represent mean ± SD fold change relative to control. P value was determined by unpaired t-test. (B) mRNA expression of NOS2 (inducible nitric oxide synthase) in BMDMs treated with control or 250 ppm CO for 24h. CO treatment significantly reduced NOS2 expression compared to control ($p = 0.003$, n = 3). Data represent mean ± SD fold change relative to control. P value was determined by unpaired t-test.

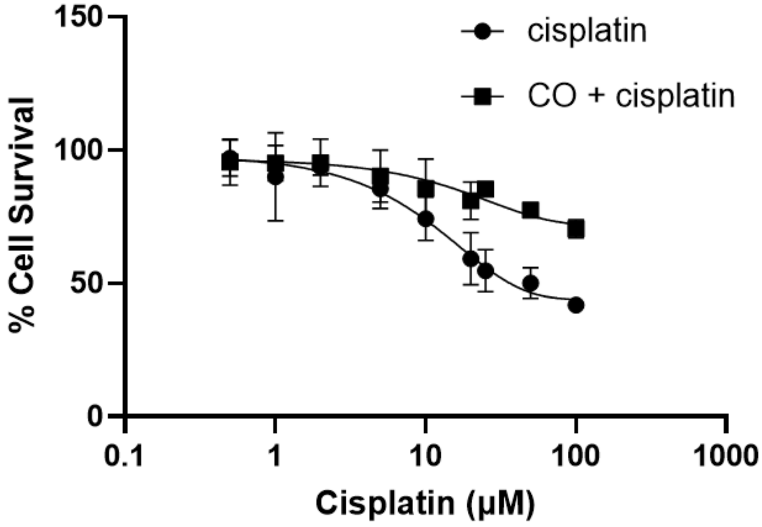


**Figure S9.** CO preserves cell viability of HK-2 renal epithelial cells upon exposure to CO (250ppm) for 3 hours + cisplatin for 24 hours (n = 3 biological replicates per group).

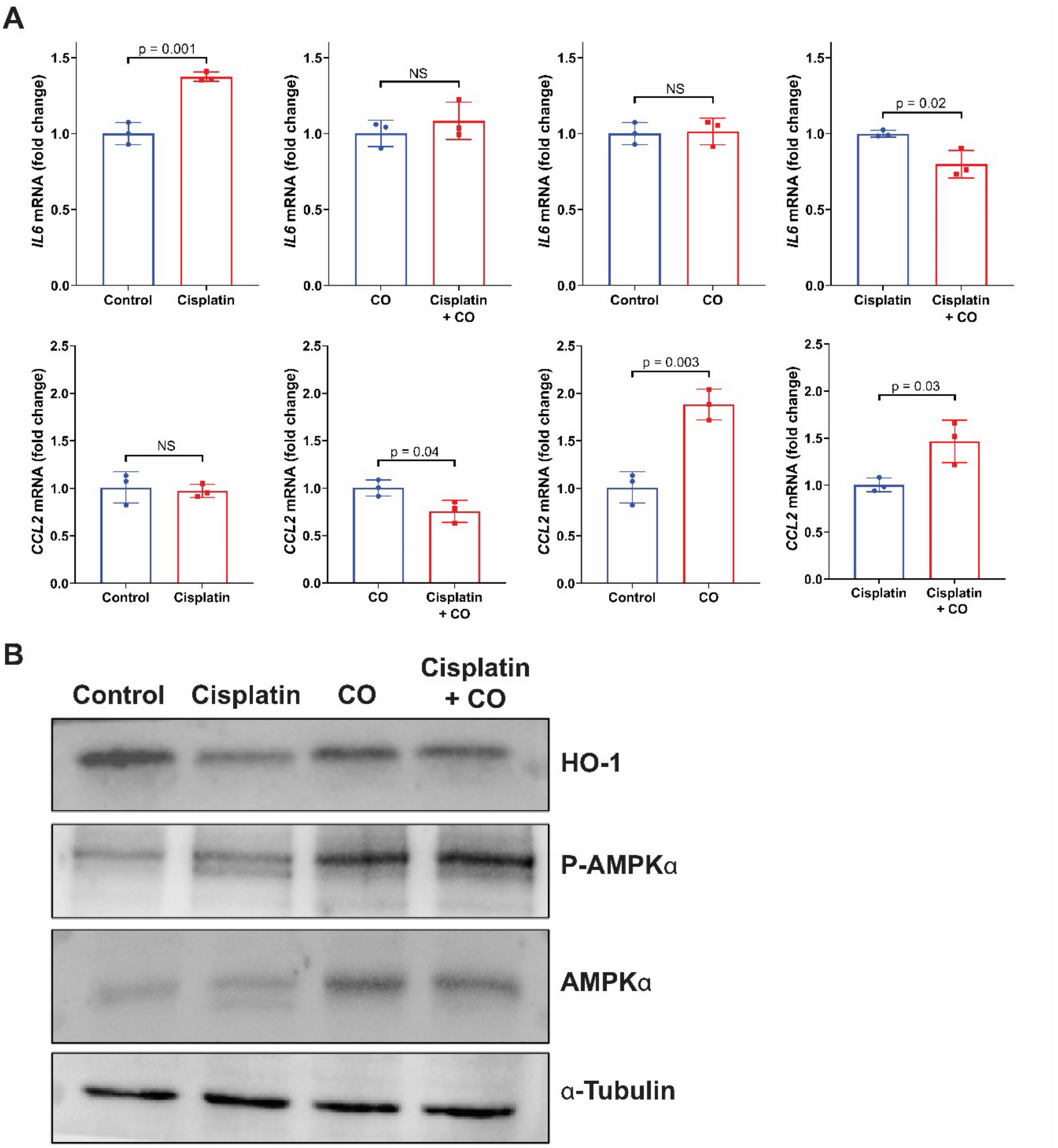


**Figure S10. CO modulates inflammatory gene expression in response to cisplatin in human normal proximal tubule epithelial cells.** (A) mRNA expression of IL6 (top row) and CCL2 (bottom row) in HK2 cells under various treatment conditions (CO:250ppm, cisplatin:10nM). Cisplatin increased IL6 expression ($p = 0.001$), which was not significantly affected by CO co-treatment (NS). CCL2 expression was not significantly changed by cisplatin alone (NS) but was reduced by CO in combination with cisplatin ($p = 0.04$). CO alone increased CCL2 expression ($p = 0.003$), and cisplatin + CO increased CCL2 compared to cisplatin alone ($p = 0.03$). Data represent mean ± SD fold change relative to control ($n = 3$). P value was determined by unpaired t-test. (B) Representative western blots showing protein levels of phosphorylated-adenosine monophosphate-activated protein kinase alpha (AMPKα), AMPKα, and α-Tubulin in cells treated with control, cisplatin, CO, or cisplatin + CO. β-actin shown as loading control.

**Table S1.** Half maximal inhibitory concentration ($IC_{50}$) upon exposure to cisplatin and 250 ppm CO. P-values were determined by unpaired T-test.

| | **$IC_{50}$ (µg/mL)** | | |
|---|---|---|---|
| **Cell line** | **Cisplatin** | **Cisplatin + CO** | **P-value** |
| OVCAR3 | 1.57 | 0.73 | < 0.0001 |
| OVCAR4 | 1.498 | 1.162 | 0.0004 |
| OV-7 | 1.38 | 0.87 | < 0.0001 |
| CAOV3 | 1.10 | 0.93 | 0.0015 |
| SKOV3 | 1.288 | 1.119 | 0.05 |

**Table S2.** Summary of HIPEC clinical trials used for ovarian cancer.

| Author or Trial Name | Registry | Trial Details | Clinical Setting | HIPEC Regimen | Primary Endpoint | Outcome |
|---|---|---|---|---|---|---|
| van Driel et al [2] | NCT00426257 | Phase 3 n = 245 | Stage III EOC, stable after 3x cycles of carboplatin/ paclitaxel, with patients set to undergo interval CRS | Cisplatin | RFS in patients receiving interval CRS with or without HIPEC | Median RFS increased to 14.2 months in the surgery plus-HIPEC group vs. 10.7 months in the group not receiving HIPEC (p = 0.003) |
| CHIPOR [6] | NCT01376752 | Phase 3 n = 415 | First relapse of EOC >6 months after completing platinum-based chemotherapy | Cisplatin | OS in patients receiving surgical resection with or without HIPEC | OS was significantly improved with addition of HIPEC (stratified hazard ratio 0.73, 95% CI 0.56–0.96; p=0.024) |
| C-HOC [7] | ChiCTR20000 28894 | Phase 2 n = 65 | Patients with EOC (FIGO stage IIIC, IVA, and IVB), unsuitable for optimal cytoreduction in primary debulking surgery | Paclitaxel | Chemotherapy response scores after IDS were compared between group that received HIPEC followed by 3x cycles of chemotherapy vs control group without HIPEC | HIPEC group exhibited a higher degree of chemotherapy response score (20.5% vs. 4.8%; p < 0.0001) as compared to control group |
| HIPECOVA [8] | NCT02681432 | Phase 3 n = 76 | Primary EOC (FIGO stages II, III, and IV) in which complete cytoreduction was achieved | Paclitaxel | OS and RFS were compared between patients receiving HIPEC with SCS vs SCS alone | Patients receiving HIPEC had improved primary outcomes (RFS: 23 months, OS: 48 months) compared to control group (RFS: 19 months, OS: 46 months) but were not significant (p = 0.22 and p = 0.579 respectively) |
| OVHIPEC [9] | NCT00426257 | Phase 3 n = 245 | Primary EOC (stage III) with no progression during at least 3x cycles of neoadjuvant carboplatin/paclitaxel | Cisplatin | PFS after undergoing interval CRS was compared between groups that received and did not receive HIPEC | Median PFS was increased to 14.3 months (95% CI 12.0–18.5) in patients receiving HIPEC as compared to 10.7 months (95% CI 9.6–12.0) in the surgery alone group (p = 0.0008) |
| Antonio et al. [10] | NCT02328716 | Phase 3 n = 71 | Primary EOC, tubal carcinoma, or primary peritoneal carcinoma (FIGO stage 3B/C) that had received 3x cycles of systemic carboplatin/ paclitaxel | Cisplatin | DFS was compared at 32 months between the experimental group receiving CRS with HIPEC and control group (CRS alone) | Median DFS was improved to 18 months in the group receiving CRS with HIPEC, as compared to 12 months in the group receiving CRS alone. HIPEC was also shown to be a protective factor against recurrence (HR = 0.12, 95 % CI 0.02–0.89; p = 0.038) |
| HORSE [11] | NCT01539785 | Phase 3 n = 167 | Primary platinum-sensitive recurrence of EOC with evidence of disease in the abdominal cavity | Cisplatin | PFS at 5 years in patients receiving HIPEC in addition to SCS vs patients receiving SCS alone | PFS at 5 years was 61.6% in the SCS group and 75.9% in the SCS plus HIPEC group (no statistical significance) |

| | | | | | | |
|---|---|---|---|---|---|---|
| Zivanovic et al. [12] | NCT01767675 | Phase 2 n = 98 | Patients with first recurrence of high-grade EOC confirmed after completion of first-line platinum-based chemotherapy and deemed resectable | Carboplatin | Proportion of patients without evidence of disease progression at 24 months following secondary cytoreduction compared between two groups (with and without HIPEC). | Median PFS was 12.3 months for patients who received HIPEC during secondary cytoreduction and 15.7 months for the group without HIPEC (p = 0.05) |
| Spiliotis et al. [13] | - | Phase 3 n = 120 | Patients with EOC (FIGO IIIC and IV) who experienced disease recurrence after initial treatment with surgery and systemic chemotherapy | Platinum-sensitive: cisplatin and paclitaxel; platinum-resistant: doxorubicin and (paclitaxel or mitomycin) | OS in patients undergoing surgical resection and systemic chemotherapy with or without HIPEC | OS was significantly prolonged at 26.7 months in the group receiving HIPEC vs. 13.4 months in the control group (p < 0.006) |
| Deraco et al. [14] | - | Phase 2 n = 26 | Treatment-naive EOC with advanced peritoneal involvement (stage III-IV) | Cisplatin and doxorubicin | OS and PFS after CRS and HIPEC | At 25 months follow-up, 5-year OS = 60.7% and 5-year PFS was 15.2% (with median = 30 months) |
| Di Giorgio et al. [15] | - | Phase 2 n = 47 | Primary advanced or recurrent ovarian cancer (TNM-FIGO stages IIIc-IV) and evidence of peritoneal carcinomatosis. All patients were set to undergo CRS + HIPEC + systemic chemotherapy | Cisplatin | OS and DFS after undergoing CRS with HIPEC and subsequent systemic chemotherapy | Mean OS was 30.4 months and mean DFS was 27.4 months. Follow up demonstrated a 5-year survival of 16.7% |
| KOV-HIPEC [16, 17] | NCT01091636 | RCT n = 184 | Patients with stage III–IV EOC who achieved optimal cytoreduction | Cisplatin | (12) PFS after undergoing CRS with/without HIPEC (13) Health-related quality of life questionnaires (3x) were assessed at baseline and after treatment in both groups. | (12) Median PFS was 18.8 months in the control group and 19.8 months in the HIPEC group (p = 0.43) (13). Health-related quality of life was maintained for those receiving HIPEC |

**Abbreviations**

PFS: progression-free survival

RFS: recurrence-free survival

DFS: disease-free survival

OS: overall survival

CRS: cytoreductive surgery

SCS: secondary cytoreductive surgery

IDS: interval debulking surgery

HGSOC: high-grade serous ovarian/fallopian tube carcinoma

EOC: epithelial ovarian cancer

** Studies that showed no statistically significant difference: Zivanovic [12], HORSE trial [11], and HIPECOVA trial [8].

** KOV-HIPEC had two publications associated with the trial [16, 17]; the first described no benefit to OS, but the second paper described quality of life outcomes to be the same in patients that received HIPEC vs. those that did not.
